# Unifying framework for the diffusion of microscopic particles in mucus

**DOI:** 10.1101/2020.07.25.221416

**Authors:** Antonio Cobarrubia, Jarod Tall, Austin Crispin-Smith, Antoni Luque

## Abstract

Mucus is a fluid that protects animals against pathogens while promoting interactions with commensal microbes. Changes in the diffusivity of particles in mucus alter viruses’ infectivity, the efficiency of bacterial pathogens to invade a host, and the effectivity of drug delivery. Multiple physicochemical properties modulate the diffusion of microscopic particles in mucus, but their combined effect is unclear. Here, we analyzed the impact of particle size, charge, chemistry, anomalous diffusion exponent, and mucus composition in the diffusivity of particles from 106 published experiments. We used a time window sampling of one second to define a consistent, effective diffusion across experiments. The effective diffusion spanned seven orders of magnitude from 10^−5^ to 10^2^ µm^2^/s. The anomalous exponent was the strongest predictor among all variables tested. It displayed an exponential relationship with the effective diffusion that explained 90% of the empirical data variance. We showed that the relationship and dominance of the anomalous diffusion exponent resulted from a general mathematical relationship obtained from first-principles for any subdiffusion mechanism. Our derivation demonstrated that the generalized diffusion coefficient is not a measurable physical quantity and must be replaced by the length and time scales associated with the underlying mobility mechanisms. This led us to a fundamental reformulation of the classic subdiffusion equation, which calls for a reinterpretation of anomalous diffusion in physical systems. We also discussed how our results impact the characterization of microscopic particle diffusion in mucus and other hydrogels.

---

Mucus is a complex fluid secreted by animals. It protects organs against the invasion of pathogens and promotes the interaction with commensal microbes (Bäckhed et al. 2005; Bakshani et al. 2018; Silveira and Rohwer 2016). Mucins—a characteristic component of mucus—are glycoproteins that form a polymeric mesh in mucus (Spagnolie 2015). Changes in the mucin network alter the diffusion of microscopic particles in mucus with disparate outcomes for the animal host. Low pH thickens mucus, reducing, for example, the diffusion and infection rate of viruses like HIV (Lai et al. 2009). Interaction with mucins also alters the diffusivity of particles in mucus. Commensal viruses that infect bacteria and reside in the gut, for instance, display immunoglobulin-like domains that are attracted to mucins, which reduces their diffusivity and increase their infectivity against bacteria (Barr et al. 2013, 2015). The regulation of particle diffusion in mucus is thus paramount for animal health, and the enhancement of diffusivity is also key for the delivery of medical drugs in the body (Cone 2009). The combined effect of these different physicochemical factors, however, remains puzzling.

Small biomelecules and biomolecular complexes tend to diffuse more readily through mucus, while larger particles are caught in the mucin network (Amsden and Turner 1999; Cone 2009). On the other hand, non-adhesive polystyrene particles with a diameter of 500 nm diffuse faster than smaller particles (200 nm) of the same type (Lai et al. 2007). Therefore, parameters other than particle size must play a role. Neutrally charged particles, for instance, display higher diffusivity than negatively and positively charged particles of the same size in mucus with a net negative charge (Abdulkarim et al. 2015; Arends et al. 2013; Hansing et al. 2016; Lieleg et al. 2010; Li et al. 2013). Increase of salt concentration shields charged particles, leading to diffusivities similar to neutrally charged particles (Arends et al. 2013; Lieleg et al. 2010; Hansing et al. 2016). Instead, low pH increases the distribution of negative charges in mucins altering the electrostatics as well as viscoelasticity of mucus, reducing diffusivity for most particles (Celli et al. 2009; Lai et al. 2009; Lieleg et al. 2010; Spagnolie 2015; Suk et al. 2011). Each of these studies show how different physical properties can controll the diffusion of microscopic particles in mucus, but the emerging picture is complex, and it is unclear if any of these physical factors is more dominant than others.

To assess the combined impact of each factor, we studied twenty-three published articles measuring the diffusion of particles in mucus or mucus-like hydrogels. Ten studies contained diffusion data that could be compared at the same time scale for spherical nanoparticles (Abdulkarim et al. 2015; Barr et al. 2015; Lai et al. 2007, 2009; Lieleg et al. 2010; Olmsted et al. 2001; Newby et al. 2017; S.Schuster et al. 2013; Suk et al. 2011; Yildiz et al. 2015). Using WebPlotDigitizer (Rohatgi 2019), we extracted 106 measurements of effective diffusion, measured at a window time of one second, that is,

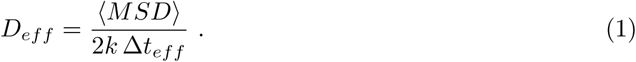

Here ⟨*MSD*⟩ was the ensamble mean squared displacement for each particle tracking experiment, *k* was the dimensions of the particle diffusivity, and Δ*t*_*eff*_ = 1 sec was the effective sampling time window (Huang et al. 2013). In all measurements, we estimated particle hydrodynamic diameter, particle type, mucus source, dominant mucin expression, and temperature. When possible, we extracted or derived the anomalous diffusion exponent (*α*), particle charge, mucus pH, mucus salt concentration, and mucin concentration. The anomalous exponent was obtained from the classic subdiffusion equation:

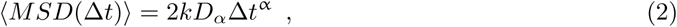

Here *D*_*α*_ is the generalized diffusion constant and Δ*t* is the sampling time window (Metzler et al. 2014).

Table 1 displays the ranges obtained for each physical parameter, and Table S.1 contains the full data set. The effective diffusions spanned seven orders of magnitude, from *∼* 10^−*2*^ µm^*2*^/s to 10^*5*^ µm^*2*^/s. Particle diameter (*d*) spanned three orders of magnitude, from 1 nm to 1,300 nm; three quarters of the particles (n=80, 75%) had diameters greater than 100 nm. The anomalous exponent (*α*) ranged from strongly subdiffusive (*α* ≈ 0.1) to purely diffusive (*α* ≈ 1), but it was obtained only for a third of the dataset (n = 33, 30%). The zeta potential (*ζ*) measured the effective surface charge of particles in solution (Kumar and Dixit 2017). The values ranged from −70 mV to +40 mV and were obtained for half of the dataset (n = 57, 52%). The temperature ranged was narrow, 295 K to 310 K. The pH ranged from mildly acidic (pH = 3.0) to slightly alkaline (pH = 7.4). Mucus experiments were associated to cell lines from four sources: human respiratory, human cervix, pig gastric, pig intestines. The dominant mucins were MUC5B, MUC2, and MUC5C. A third of the experiments were conducted in artificial mucus-like hydrogels.

**Table 1:**
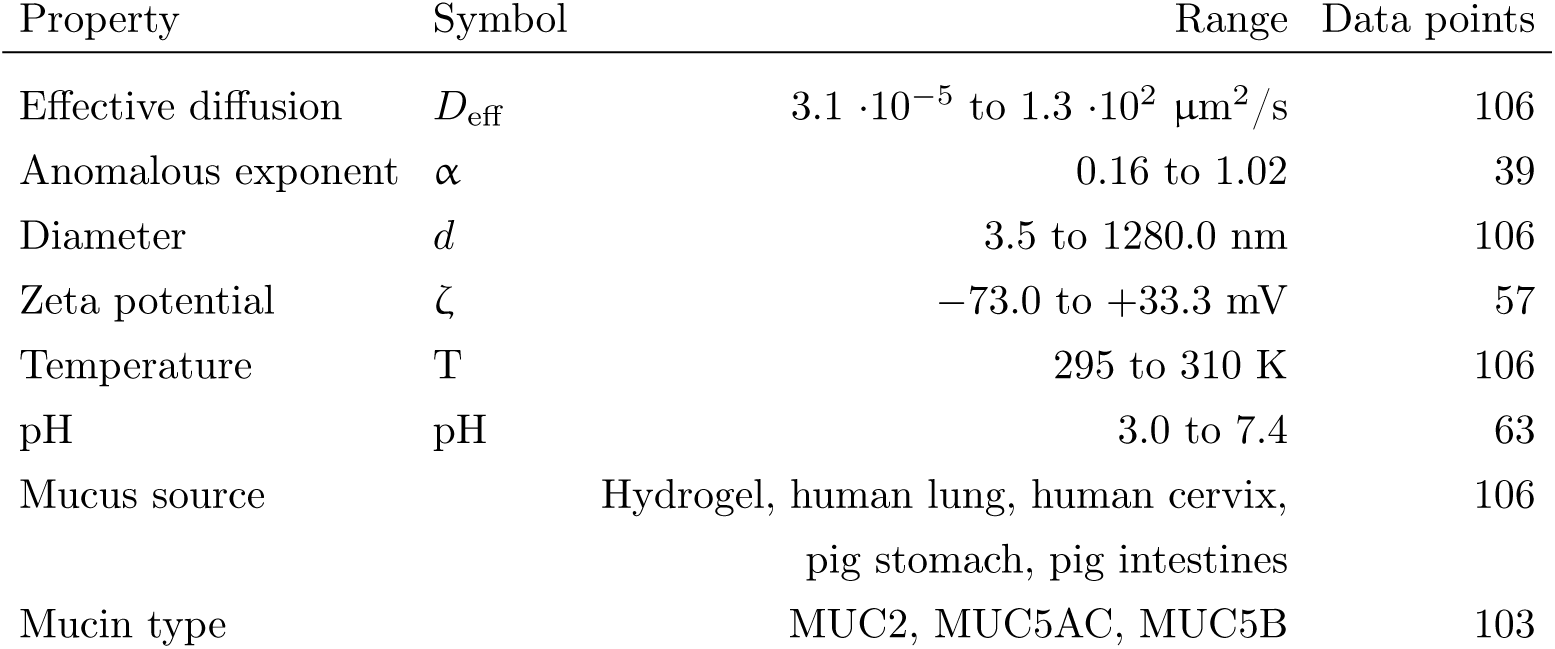
Summary of empirical data. *D*_*eff*_ : The effective diffusion was obtained for a common time window of 1 second. For references that shared relative diffusion with respect diffusion in water, the effective diffusion constant was scaled using the Stokes-Einstein equation using the temperature and hydrodynamic particle diameter reported (Miller 1924). Room temperature (298 K) was assumed if temperature was not reported in the study. Particle type data was obtained by classifying particles as COOH, PEG, virus, amine, antibody/protein, or chitosan. This was chose as a qualitative measure of particle-mucin bonds. The dominant mucin composition from each mucus source was obtained by evaluating the expression levels of mucin genes from the genome bioinformatics portal Ensembl (Zerbino et al. 2018). Mucins were identified assuming the tissue/organ associated with each mucus, or closely associated tissues. Expression levels were collected by taking the average of reported median of transcript per million (TPM) RNA-sequence and the most explicitly stated expression levels of low, medium, and high. Based on potential gene expression of mucins with reported levels of below cutoff, TPM measured below the minimum (0.05 TPM) is distinguished from experiments with no data due to possible gene expression. An expression level of low, medium or high was obtained over reports of below cutoff in the same tissue. The dominant mucin was determined by the highest expression level then, if necessary, by the highest average of median TPM. Identification of mucin expression based on tissues was associated with each mucus: human respiratory mucus and human cystic fibrosis mucus were associated with the human lung mucin genes; human cervical mucus and cervicovaginal mucus were associated with human cervix or uterus mucin genes; pig intestinal mucus was originally from jejunum part of the small intestine, however, due to a lack of reports for jejunum tissue, the associated mucin genes were taken as the average of the median of TPM of pig duodenum and pig ileum parts of the small intestine based on the close proximity to the jejunum; pig ileum intestinal mucus were associated with ileum tissue mucin genes; pig gastric mucus were collected from pig stomach mucin genes.

The non-parametric statistical method random forest was applied to identify the most relevant variables impacting the effective diffusion. The variables pH, mucus concentration, salt concentration, and MUC5B were omitted because they were missing for most data in the multivariate analysis. The selection of variables was obtained in two rounds, discarding not statistically significant variables (p-value > 0.05) in each round. This led to five significant variables (Figure 1). The anomalous diffusion exponent (*α*) was the most dominant variable with an average percentage increase in mean square error (%MSE) of 22.4(±3.2)% (S.D.) (p-value = 0.0099). The second most dominant variable was particle type, followed by zeta potential, mucus source, and dominant mucin.

**Figure 1:**
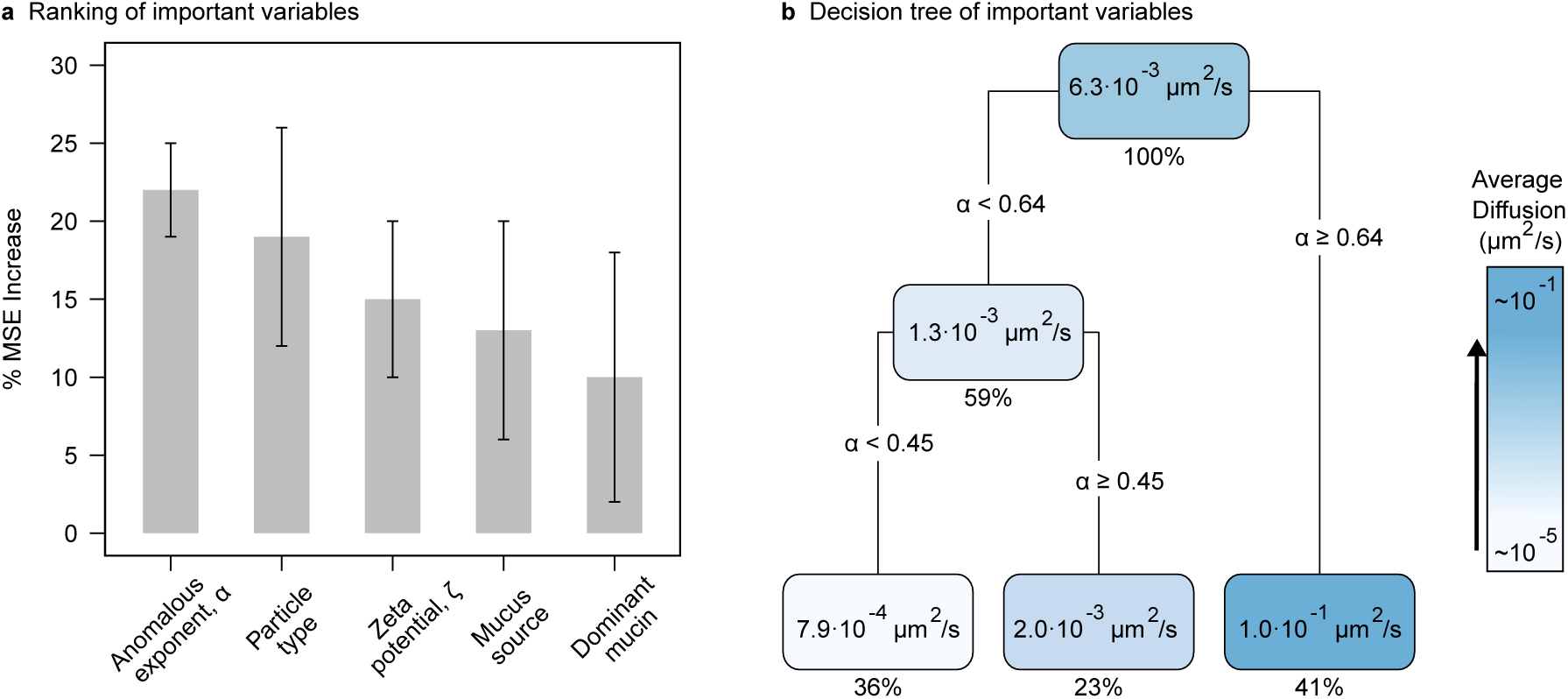
Selected variables impacting effective diffusion. **a**, Average percentage increase of mean-square error (% MSE) for the selected variables. These variables were investigated in permutations of three using random forest (R package rfPermute by Archer 2019). The error bars correspond to the standard deviation. **b**, Decision tree for the most important variables. Each node contains the predicted average *D*_eff_ and percentage of data predicted. The gradient display diffusion values from *∼* 10^−5^ µm^2^/s (white) to *∼* 10^−1^ µm^2^/s (blue).

When analyzing the selected variables individually, the anomalous diffusion exponent displayed by far the strongest correlation with the effective diffusion (non-parametric Spearman correlation *ρ* = 0.93, p < 2.2E-16***, n = 39). The effective diffusion increased exponentially with the anomalous diffusion exponent (Figure 2a). The linear regression for the transformed data (log-linear) explained 89% of the variance (slope=5.3 ± 0.3, p-value *<* 2.0 · 10^−16⋆ ⋆ ⋆,⋆^ *R*^2^ = 0.89, least-squares method). The anomalous exponent, however, was only reported for *∼* 37% (n=33) of the data, which included carboxylated, pegylated, and viral particles. An inverse statistical model was fitted to estimate the mean anomalous exponents as a function of the effective diffusions for the remaning 63% of the data, corresponding to amine, chitosan, antibodies, and proteins particles (n = 67). Particles with effective diffusions above *D*_*eff*_ *>* 10 µm^2^/s were predicted to display regular diffusion (*α ∼* 1).

**Figure 2:**
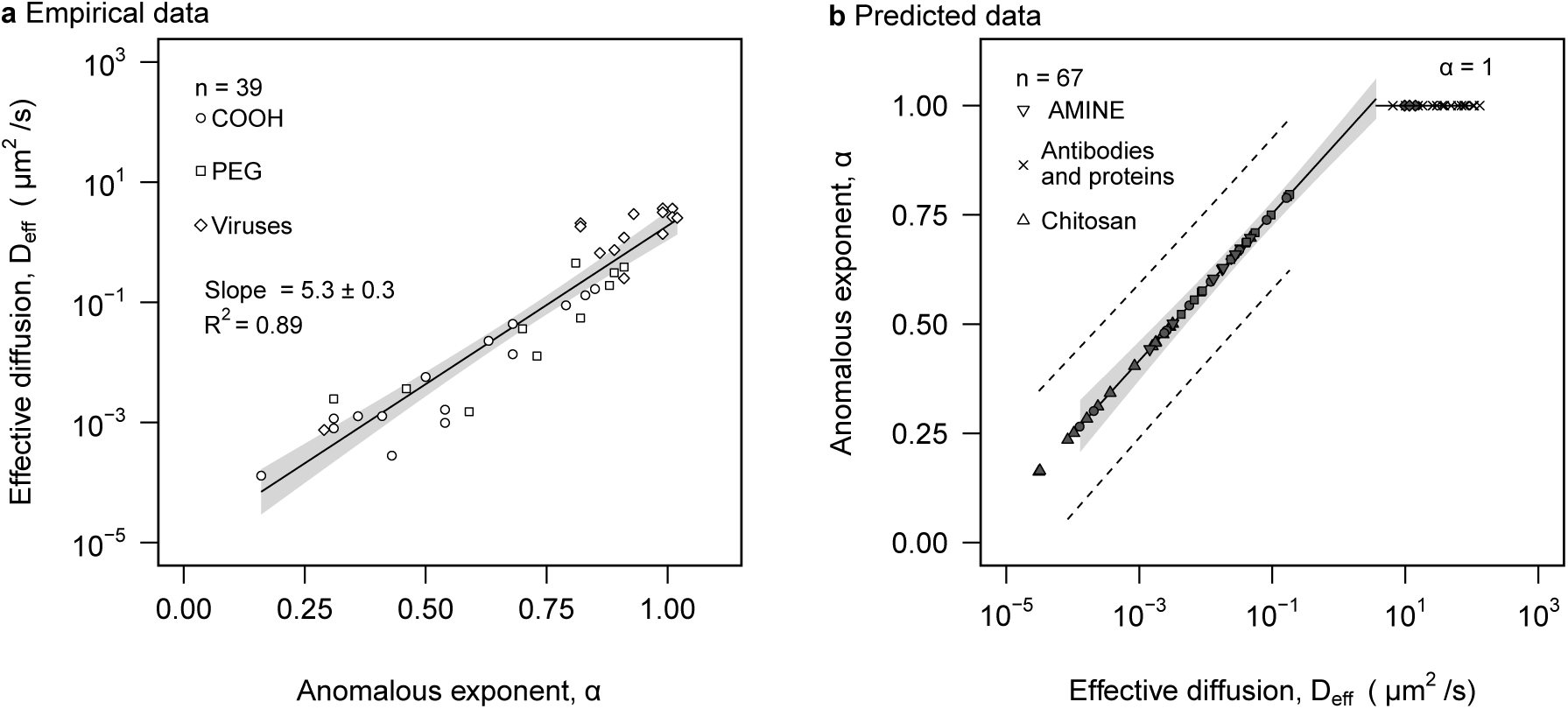
Effective diffusion and anomalous exponent analysis. **a**, effective diffusion was plotted as a function of anomalous exponent. The solid line represents the regression model. The grey area represents the 95% confidence interval. Statistically significant slope and *R*^2^ of linear regression is displayed. **b**, anomalous exponent was predicted based on the model found empirically in **a**. The solid line designates the predicted linear model. The grey area represents the 95% confidence interval of the predicted linear model. The dashed line represents a 95% prediction interval **a-b**, distinguished particle types are represented in the legend of both panels..

To elucidate the physical origin of the dominance of the anomalous exponent (*α*), its relationship with the effective diffusion, *D*_*eff*_, was derived from Eqs. (1) and (2):

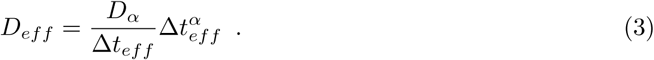

The effective diffusion, thus, has a factor that varies exponentially with the anomalous exponent (*α*). However, it also depends on the generalized diffusion (*D*_*α*_), which is implicitly a function of the anomalous diffusion exponent as well as particle and fluid properties, and its functional form changes depending on the specific underlying subdiffusion mechanism (Metzler et al. 2014; Joiner et al. 2019). Our meta-analysis contained a broad range of data (Table 1), including particles with different chemistry, mucus of different types, different physicochemical conditions, and independent groups carrying different experimental implementations. Therefore, it was not obvious how the generalized diffusion would be changing in each case, and Eq. (3) was not sufficient to justify the dependence and dominance of *α* in determining the mobility of particles in mucus. To understand this phenomenon, the generalized diffusion had to be analyzed further.

The units of the generalized diffusion constant, *D*_*α*_, depend on the anomalous exponent. In our study, these units were µ*m*^*2*^*/s*^*α*^. The anomalous diffusion exponent, as any other physical quantity, has an associated uncertainty (error or standard deviation) (Taylor 1997). Thus, the units of *D*_*α*_ are uncertain. In other words, the generalized diffusion constant is not measurable. The fact that *D*_*α*_ is not a physical quantity has been previously overlooked and mandates a revision of the classic subdiffusion equation, Eq. (2).

To reformulate the subdiffusion equation, we split the generalized diffusion into the characteristic length scale (*L*_*D*_) and characteristic time scale (*t*_*D*_) associated to the physical mechanism responsible for the mobility of the particle:

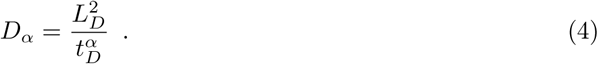

This ansatz was combined with the classic subdiffusion equation, Eq. (2), obtaining:

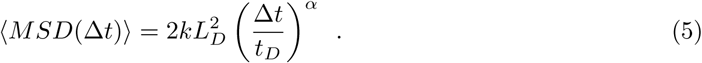

This reformulated subdiffusion equation is valid for window times larger than the characteristic mobility time scale, Δ*t ≥ t*_*D*_. For smaller window times, the underlying mobility mechanism will dominate, requiring a different formulation for the displacement (Joiner et al. 2019).

The reformulated subdiffusion equation, Eq. (5), was combined with the definition of the effective diffusion, Eq. (1), obtaining

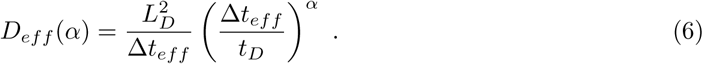

The effective diffusion, thus, depends exponentially on the anomalous diffusion exponent, *α*, explaining the relationship observed empirically for the effective diffusion of particles in mucus (Figure 2). To investigate the origin of the dominance of the anomalous diffusion exponent in the variation of the effective diffusion across multiple scales, we investigated the logarithm of the effective diffusion equation, Eq. (3):

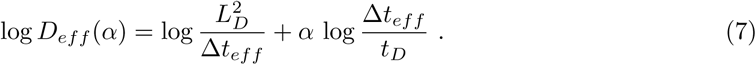

For a fix time window, Δ*t*_*eff*_, the change of the effective diffusion with respect the anomalous diffusion is *∂* log *D*_*eff*_ */∂α* = log (Δ*t*_*e*_*ff/t*_*D*_), while the the impact of the change in the characteristic mobility length and time scales are, respectively, *∂* log *D*_*eff*_ */∂* log *L*_*D*_ = 2 and *∂* log *D*_*eff*_ */∂* log *t*_*D*_ = −*α*. The changes were evaluated with respect the logarithms of the length and time scales to obtain results independent of the measuring units. The change with respect the length scale is constant with a value of two, while the change with resepct the time scale is bounded within an absolute value of one. Thus, for sampling time windows that are more than two orders of magnitude larger than the characteristic mobility time scale, Δ*t*_*eff*_ */t*_*D*_ *»* 10^2^, the anomalous diffusion would be the dominant physical factor determining the change in the effective diffusion.

This hypothesis was confirmed for the collected diffusion data in mucus, Eq. (7), by extracting the average mobility length and time scales from the empirical data (Figure 2**a**). This led to *L*_*D*_ *∼* 3 nm and *T ∼* 5 µs. The sampling window time was Δ*t*_*eff*_ = 1 s. Therefore, Δ*t*_*eff*_ */t*_*D*_ *∼* 10^6^ *»* 10^2^, satisfying the condition for the dominance of the anomalous diffusion exponent derived above. To justify the values obtained for the average length, *L*_*D*_ and time scales *t*_*D*_, it was necessary to look into the underlying mechanisms fueling the mobility of the particles. A given mechanism would propel the particles with a velocity *v*_*D*_ for the characteristic time *t*_*D*_. This defines the characteristic length scale *L*_*D*_ *∼ v*_*D*_*t*_*D*_. In all experiments analyzed, the particles were passive, acquiring a transient velocity fueled by the transfer of kinetic energy from the thermal buffeting of the fluid, that is, 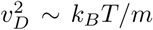, where *k*_*B*_ is the Boltzmann constant, and *m* is the mass of the particle. Mucus is a viscous fluid, and this velocity will dissipate with a characteristic time *t*_*D*_ *∼ m/γ*, where *γ* is the friction coefficient. Not surprisingly, this leads to the Stokes-Einstein equation for the underlying characteristic diffusion, 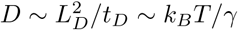. For the typical mid-size particle in the data analyzed, *d ∼* 100 nm, the relaxation time is *t*_*D*_ *∼* 1 µs and the microscopic diffusion is *D ∼* 1 µ^2^/s. This leads to the characteristic mobility length scale 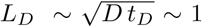 nm (Joiner et al. 2019). Therefore, the estimated values for *L*_*D*_ and *t*_*D*_ are consistent with the average empirical values obtained for Eq. (7), supporting our reformulated subdiffusion framework. Thus, the problem of characterizing the diffusion of a particle in mucus reduces to identifying the physical factors that regulating the anomalous exponent. These factors depend on the specific mechanism hindering the regular diffusion (Metzler et al. 2014). There are at least two mechanism that may play an important role in mucus. First, microscopic particles can bind to the mucin fibers that constitute mucus leading to subdiffusion (Barr et al. 2015). Second, mucin fibers form a polymeric mesh that can trap particles as observed in other hydrogels (Wong et al. 2004). Below we discuss the physical factors that control the subdiffusion exponent in each case.

Binding to mucins in mucus does not necessarily lead to subdiffusion. If a particle has a single binding site with a characteristic binding time *t*_*b*_, this will elongate the characteristic diffusion time, *t*_*D*_ *∼ t*_*r*_ + *t*_*b*_ leading to the microscopic diffusion 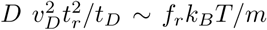. Therefore, the diffusion will be rescaled by the fraction of time spent in the relaxation of the thermal energy, *f*_*r*_ = *t*_*r*_*/*(*t*_*r*_ + *t*_*b*_), without altering the anomalous exponent. However, if more than one region of the particle can bind to mucins simultaneously and the number of number of regions bound to mucines vary stochastically, an increase of the binding time beyond the sampling time, *t*_*b*_ *»* Δ*t*_*eff*_, would lead to an effective power law distribution of binding times with no apparent characteristic binding time Xu et al. 2011. The emergence of long-tailed attachment time distributions leads to subdiffusion. The anomalous exponent, *α*, is equal to the exponent, *ν*, of the asymptotic approximated power-law distribution of attachment times (Metzler et al. 2014; Joiner et al. 2019). The generalized subdiffusion constant is given by

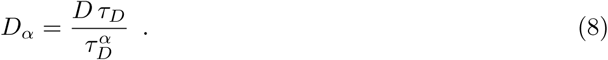

This was obtained using a continuous-time random walk approximation (Joiner et al. 2019). Here, *D* is the diffusion of the particle in the absence of interactions with mucins, *tau*_*D*_ is the average diffusion time of a particle before attaching again to a mucin fiber. This result is consistent with the ansatz that we introduced in Eq. (4). The anomalous exponent can also be related to the average minimum time of a particle attached to a mucin fiber (*τ*_0_):

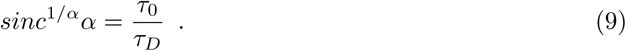

This mechanism indicates that particle-mucin affinity will dominate the effecitive diffusion of a particle in mucus. Unfortunately, the experiments analyzed did not explored the particle affinities to mucus explicitely.

The microenvironment trapping mechanism was observed in F-actin networks, where microscopic tracers were shown to follow anomalous diffusion. The anomalous exponent was a linear function of the ratio between the particle size (*d*) and network’s mesh size (*ξ*) (Wong et al. 2004). The empirical dependency obtained was *α* ≈ 1 for *d/ξ <* 0.1, *α* ≈ −1.25 *d/ξ* + 1.38 for 0.1 *< d/ξ <* 1.1, and *α* ≈ 0.1 for *d/ξ >* 1.1. Thus, particles with a size that is 10% of the mesh size or smaller diffused normally, while particles with a size similar or larger to the mesh or displayed a reduced diffusitivity with a low anomalous exponent. This phenomenon was justified qualitatively assuming an elastic energy threshold that is eventually large enough to overcome the free energy barrier and push the particle over a new microenvironment. The specific parameters of the relationship were not derived from first principles, but one would expect a similar behavior in mucus. This mechanism indicates that the effective diffusion of relatively large particles will be severely affected independently on particle-mucin interactions, that is, the particle chemistry. Unfortunately, the mesh size was not measured or reported in most experiments reviewed in our study.

Particle size was not an apparent significant predictor in the random forest analysis (Figure 1**a**), but the subdiffusion mechanisms discussed above indicated that it should be relevant when approaching the typical mesh size of mucus. The analysis of the effective diffusion as a function of particle diameters indicated a clear threshold around *d ∼* 100 nm (Figure 3**a**). Larger particles, *d >* 100 nm, displayed lower effective diffusion values but with no apparent statistical correlation with size (*ρ* = −0.24, p= 0.19). The associated empirical and predicted anomalous exponents ranged from 0.15 to 1, indicating that factors other than particle size are influencing the subdiffusion. Smaller particles, *d <* 100 nm, displayed an effective diffusion with a significant statistical correlation (Figure 3**a**). In particular, those particles that had been predicted to display regular diffusion were inversely dependent with particle size, that is, slope *m ∼* −1 (Figure 3). As predicted by the microenvironment trapping mechanism, small particles in mucus displayed regular diffusion, midsize particles were subject to subdiffusion (although the attachment-mechanism cannot be discarded), and large particles display a variety of outputs probably dependent on the mesh size (and potential interaction with mucus). The average mucus in humans has a typical mesh size between 100 to 1000 nm (Cone 2009), which explain the diffusion behavior for particles around 100 nm, or greater, in Figure 3**b**.

**Figure 3:**
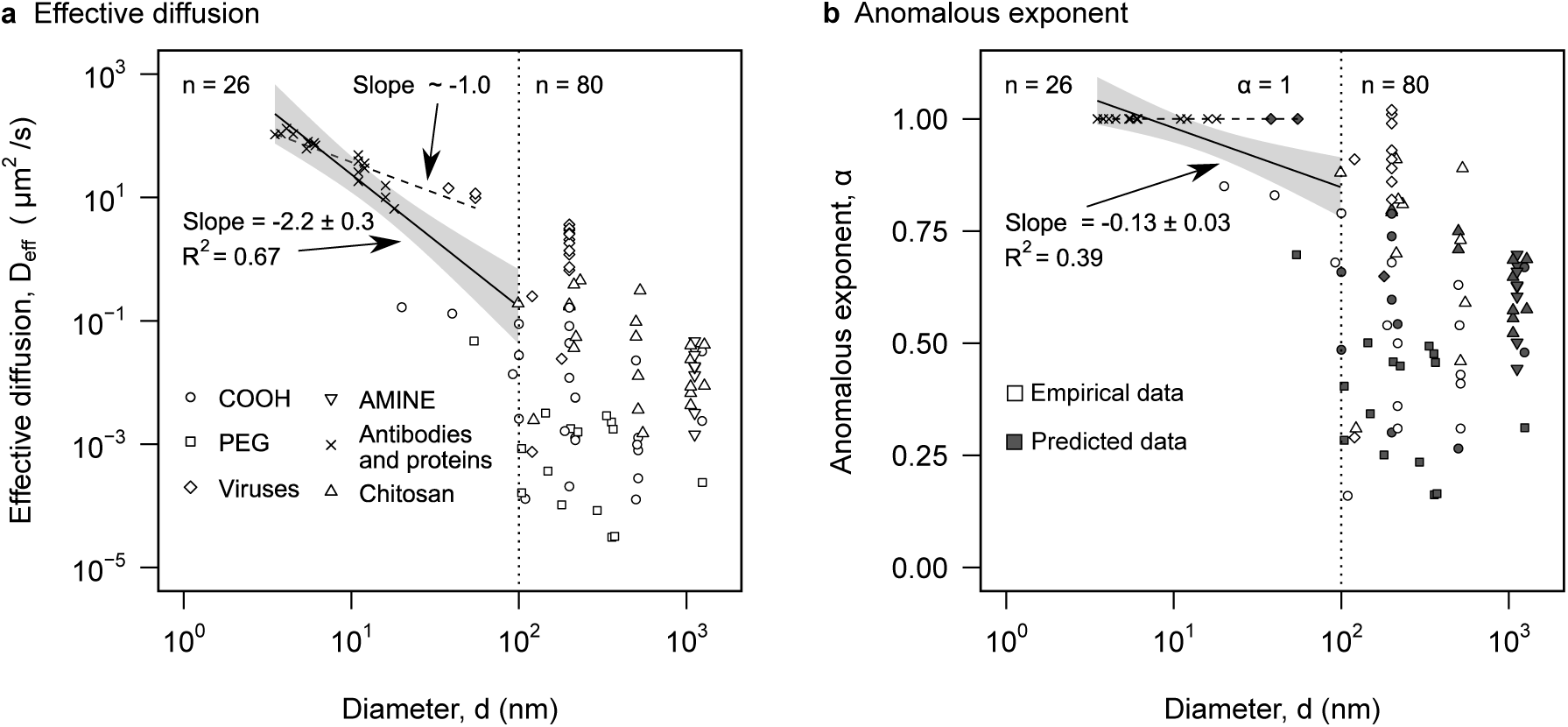
Particle size analysis. **a**, effective diffusion was plotted against particle size, d. **b**, anomalous exponent was plotted as a function of particle size. **a-b**, different particle types with different symbols are indicated in **a**’s legend. Separate analysis was conducted for particles smaller than 100 nm represented by the dotted line at d = 100 nm. Solid line corresponds to significant linear regression and the grey area represents the 95% confidence interval. Significant slope and *R*^2^ of each linear regression are displayed.

Particle type was selected as the second most relevant variable to predict the effective diffusion based on the random Forest analysis (Figure 1). Comparing the effective diffusion for the different particles confirms this prediction (Figure S.1**a**). Antibodies and proteins displayed the fastest effective diffusion with a mean of 48.9 µm^2^/s (Figure S.1). Viruses were the second fastest group with a mean effective diffusion an order of magnitude smaller, 3.5 µm^2^/s. Pegylated and amine particles formed the third group. They displayed statistically similar effective diffusions with means (medians) 0.99 µm^2^/s and 2 · 10^−2^ µm^2^/s. This was followed by COOH particles, mean (median) 3 · 10^−2^ µm^2^/s, and finally chitosan 4 · 10^−3^ µm^2^/s. Difference in particle size could explain the reduction in effective diffusion for antibodies/proteins, viruses, and PEG particles (Figure S.1**b**). They had, respectively, median sizes of *∼* 10 nm, *∼* 100 nm, and *∼* 1000 nm. It is unclear what were the physico-chemical factors behind the slower diffusion of Amine, COOH, and Chitosan particles (Figure S.1).

The third predictor for effective diffusion was particle charge, express as the zeta-potential (Figure 1). Particles with negative zeta potential displayed a positive correlation with the effective diffusion constant with a Spearman correlation of *ρ* = 0.6 (p = 0.002***, n = 36) (Figure 4**a**). The relationship was approximated by an exponential function, *D*_*eff*_ *∼ exp*(*mξ*). The potential rate, *m*, was *m* = 0.024 ± 0.006 (p *∼* 0.0002***) obtained from a least-square linear regression using the log-linear data. This exponential model accounted for 30 % of the variance (*R*^2^ = 0.30). The largest effective diffusions were achieved at neutral zeta potentials. Positive zeta potentials (n=21) had lower values but did not display a statistically significant correlation the effective diffusion. Particle size or other properties did not seem to explain the trend observed for negatively charged zeta potentials. These particles, however, displayed a linear positive correlation with the anomalous diffusion (Figure 4**b**). Based on the two mechanisms discussed above, one interpretation could be that given the negative charge of the mucin fibers, an increasing negative charge of a partice will increase its effective radius, increasing the particle size to network mesh ratio and thus reducing the diffusivity. Alternatively, the presence of negative charges could compete for ions with respect the mucin fibers exposing hydrophobic regions that could interact with the particles as has been observed in carboxilated particles forming bundles with mucus (Lai et al. 2007, 2009). Based on the attachment-mechanism, the increase in negative zeta potential would be proportional to the attachment time induced. The positive zeta potential would be expected to interact with mucin fibers. That could explain the reduction inf effective diffusion with respect neutral structures, but it did not display any apparent correlation with the anomalous exponent.

**Figure 4:**
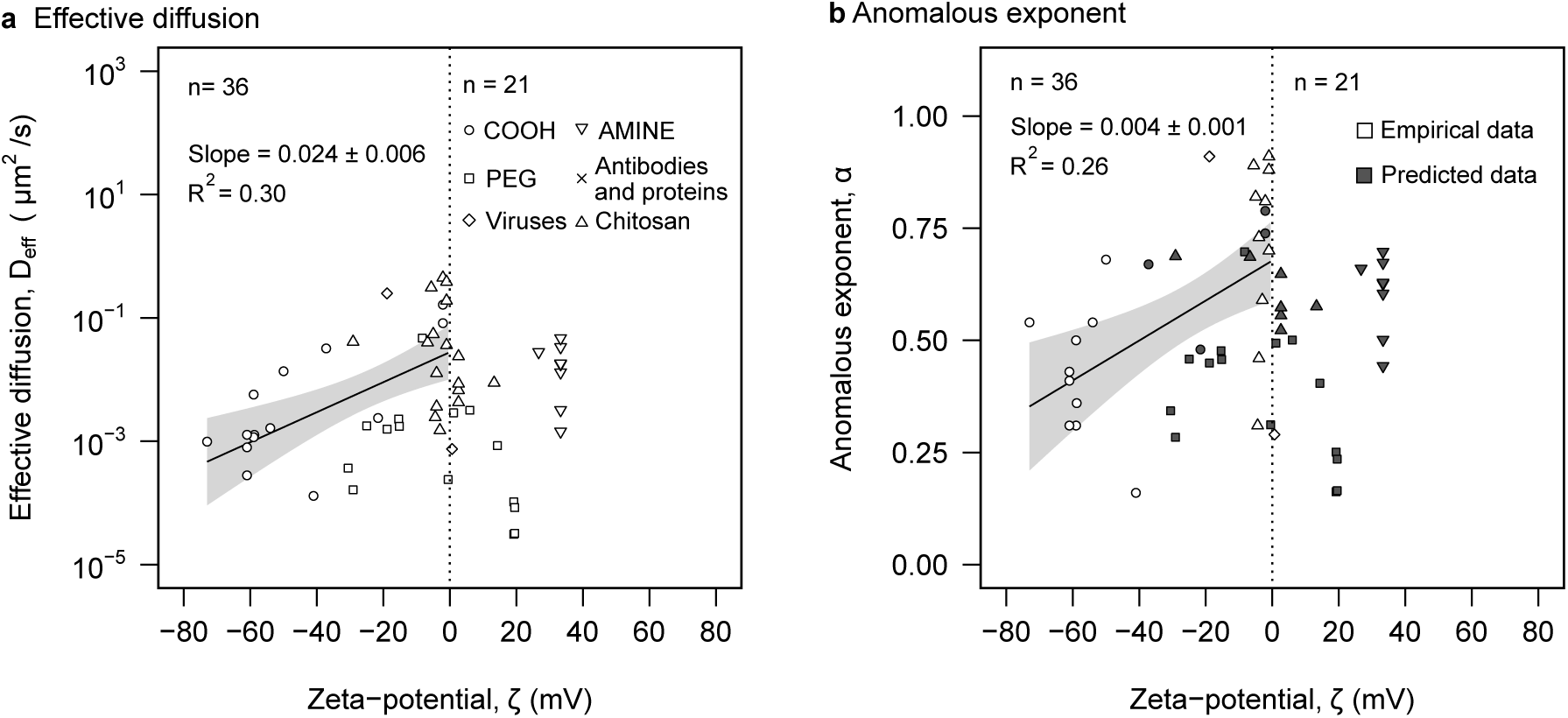
Electrostatic analysis. **a**, effective diffusion was plotted against zeta potential. **b**, anomalous exponent as a function of zeta potential. Classification of empirical and predicted data is represented in the legend. **a-b**, Different particle types are distinguished in **a** legend. Separate tests based on charge sign was conducted and designated by the dotted line at *ζ* = 0. Solid line represents significant linear regression. Grey area represents the 95% confidence interval of the linear regression. Significant slope and *R*^2^ of each linear regression are displayed.

The mucus source and dominant mucin were the last two significant predictor of effective diffusion. The effective diffusion was faster in human cervix samples with a median *∼* 10 µm^2^/s, although the values spanned six orders of magnitude, from *∼* 10^−4^ to *∼* 10^2^ µm^2^/s (Figure S.1**a**). The effective diffusion was the slowest in mucus from human lung (median *∼* 10^−2^ µ^2^/s) and pig intestine (median *∼* 10^−*2*^ µm^*2*^/s). The median particle size in empirical data from human cervix mucus was more than an order of magnitude smaller, *∼* 10 nm, than for the empirical data from the other sources. The median pH for the empirical data from human cervix mucus was significantly lower pHs (median 5.5) compared to the other sources (median 7). Lower pH tends to thicken mucus (Hwang et al Rheological Properties of Mucus 1969), thus expecting a slower effective diffusion. But the particle size may have offset this trend. The transcription analysis identified MUC5B, which is dominant in human cervix, displaying the largest effective diffusion (median *∼* 10 µm^2^/s) compared to the other dominant mucins, MUC2 common in respiratory mucus (median diffusion *∼* 10^−1^ µm^2^/s), and MUC5AC common in intestinal mucus (median diffusion *∼* 10^−2^ µm^2^/s) (Figure S.3).

Thus, our meta-analysis discovered that the anomalous exponent is the dominant factor regulating the effective diffusion microscopic particles in mucus, explaining 90% of the variance across 6 orders of magnitude (Figure 2**a**). However, less than 40% of the empirical data had measured the anomalous exponent, indicating an important gap in the field about the dominance of this factor in the diffusion of particles in mucus. To empirically validate our finding, we provide predictions of the anomalous exponent in Figure 2**b**. Remarkably, those particles predicted to display regular diffusion showed an inverse dependence between the effective diffusion and particle size, as expected from standard Brownian motion (Figure 3).

Our first-principles analysis of subdiffusion provided a general equation that explained the exponential relationship between anomalous diffusion and the effective diffusion as well as the dominance of the anomalous exponent, independently of the underlying subdiffusion mechanism, Eq. (6). Our analysis indicated that the generalized diffusion constant is not a well-defined physical quantity, and it must be replaced by characteristic length and time scales associated to the underlying mobility mechanisms, Eq. (4). This led to a reformulation of the subdiffusion equation, Eq. (5), which applies to any physical system. We conclude that the physical factors regulating the anomalous exponent is key to characterize and control the diffusivity of particles. In mucus and hydrogels, in particular, we propose that the attachment and microenvironment mechanisms should be used as a guide. Measuring the attachment time distributions and mucus mesh size are key factors regulating the anomalous exponent in these mechanisms. But they had not been measured even among the mucus experiments that measured anomalous exponents. Our work, thus, fills a gap that would guide a more effective and insightful study of effective diffusions in mucus and other complex fluids. Our study also mandates a reinterpretation of the generalized diffusion constant in any physical system.

—

## Supplementary material

**Table S.1:**
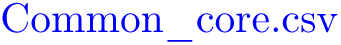
Supplementary material of common core file of all reviewed data. Common_core. The first row is a header with designated column names fitted to physical properties collected. Particle name, particle type, zeta potential, particle size, effecitve diffusion constant at 1s, anomalous exponent, diffusion in water, ratio between effective diffusion constant at 1s and diffusion in water, temperature, pH, dosing medium, salt type used, salt concentration, mucus used in the experiment, mucus concentration, mucus purification, mucin gene expression level, and dominant mucin gene is denoted as Particle, Surface_Chemistry, Zeta, Diameter, D_w, Diffusion_constant, alpha, Ratio_Diffusion, Temperature, pH, Dosing Medium, Salt_type, Salt_Concentration, Mucus_Type, Muc_Con, Purification, ‘mucin gene name’_EL, and Dominate_Mucin, respectively.

**Table S.2:**
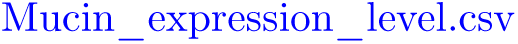
Supplementary material on dominant mucin classification based on mucus source. Mucin_expression_level. The first row is a header and first column designates mucus type. Represented expression levels are denoted as ‘mucin gene name’_EL. Expression levels are classified as below cutoff, low, medium, high, or unavailable. The average median Transcripts per million (TPM) for each mucin gene is designated as ‘mucin gene name’_TPM_median_avg. See methods for more information.

**Table S.3:** Supplementary material of linear Analysis of Nanoparticles’ Mobility Through Mucus and Biohydrogels. ‖ Simple Linear Regression and Pearson’s p-value for the slope. ‡ Spearman analysis. ¶ Effective Diffusion coefficient (*µm*^2^*/*s). ∧ Logarithmic of base 10 (*log*_10_). ◦ Diameter less than 100 nm. *** Data has strong significance. *†*Zeta potential (mV). Overview table of values associating with statistically significant simple linear regression along with spearson and pearson analysis.

| Dependent<br>variable | Independent<br>variable | Slope | Intercept | $R^2$ | p-value\|\| | rho‡ | p-value‡ |
| --- | --- | --- | --- | --- | --- | --- | --- |
| Effective Diffusion¶^ | Diameter^o | $-2.1 \pm 0.3$ | $3.5 \pm 0.4$ | 0.67 | 3.5E-7*** | -0.9 | 1.7E-12*** |
| Effective Diffusion¶^ | Negative† | $0.024 \pm 0.006$ | $-1.6 \pm 0.2$ | 0.30 | 0.0006*** | 0.6 | 0.0002*** |
| Effective Diffusion¶^ | Positive† | $0.01 \pm 0.02$ | $-2.8 \pm 0.4$ | 0.03 | 0.5 | 0.3 | 0.2 |
| Effective Diffusion¶^ | Alpha | $5.3 \pm 0.3$ | $-5.0 \pm 0.2$ | 0.89 | <2.0E-16*** | 0.9 | <2.2E-16*** |
| Anomalous Exponent | Diameter^o | $-0.13 \pm 0.03$ | $1.11 \pm 0.04$ | 0.39 | 0.0007*** | -0.6 | 0.001 |
| Anomalous Exponent | Negative† | $0.004 \pm 0.001$ | $0.68 \pm 0.04$ | 0.26 | 0.002*** | 0.5 | 0.0007*** |
| Anomalous Exponent | Positive† | $0.003 \pm 0.003$ | $0.44 \pm 0.07$ | 0.04 | 0.4 | 0.3 | 0.2 |

**Figure S.1:**
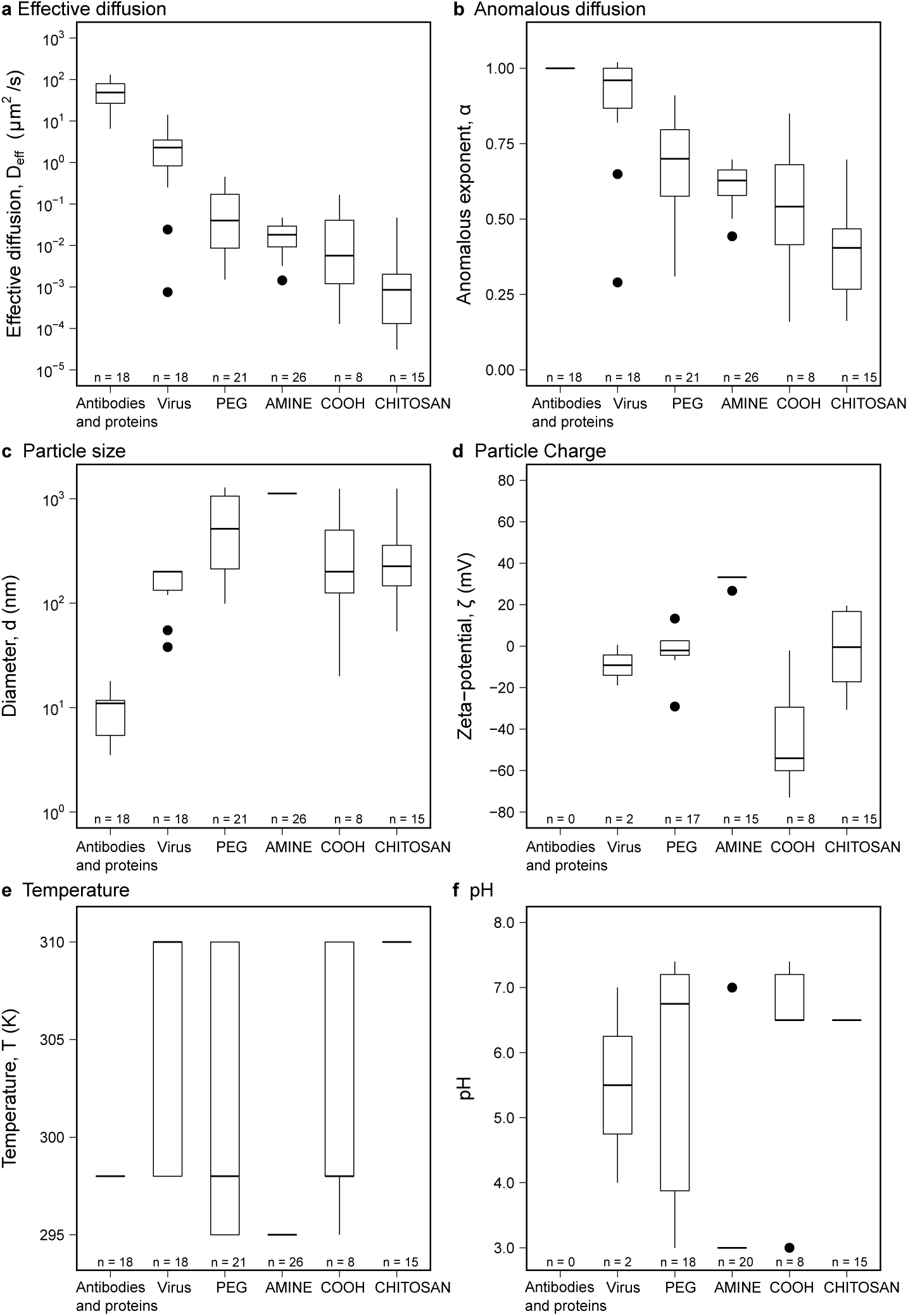
Transport capabilities based on particle type. **a**, effective diffusion constant at one second based on particle type **b**, anomalous exponent based on particle type. **c**, .particle size based on particle type. **d**, particle net charge based on particle type. **e**, mucus temperature based on particle type. **f**, mucus pH based on particle type. **a-f**, box plots are ranked by effective diffusion from high to low. The total amount of data points for each particle type is designated as n aligned with their respected particle type for each panel.

**Figure S.2:**
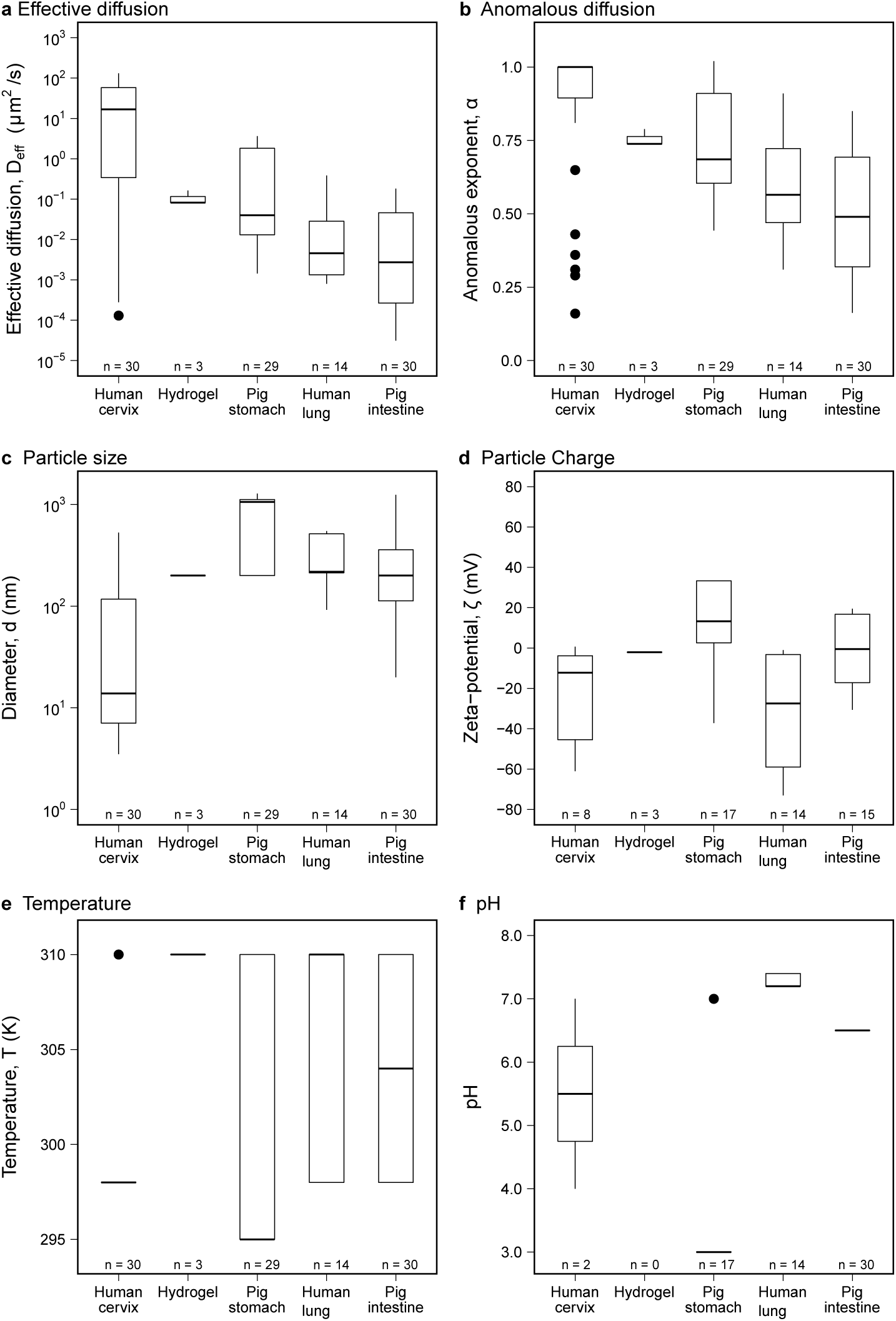
Mucus source influence on particle transport. **a**, effective diffusion constant at one second based on mucus source. **b**, anomalous exponent based on mucus source. **c**, particle size based on mucus source. **d**, particle net charge based on mucus source. **e**, mucus temperature based on mucus source. **f**, mucus pH based on mucus source. **a-f**, box plots are ranked by effective diffusion from high to low. The total amount of data points for each mucus source is designated as n aligned with their respected mucus source for each panel.

**Figure S.3:**
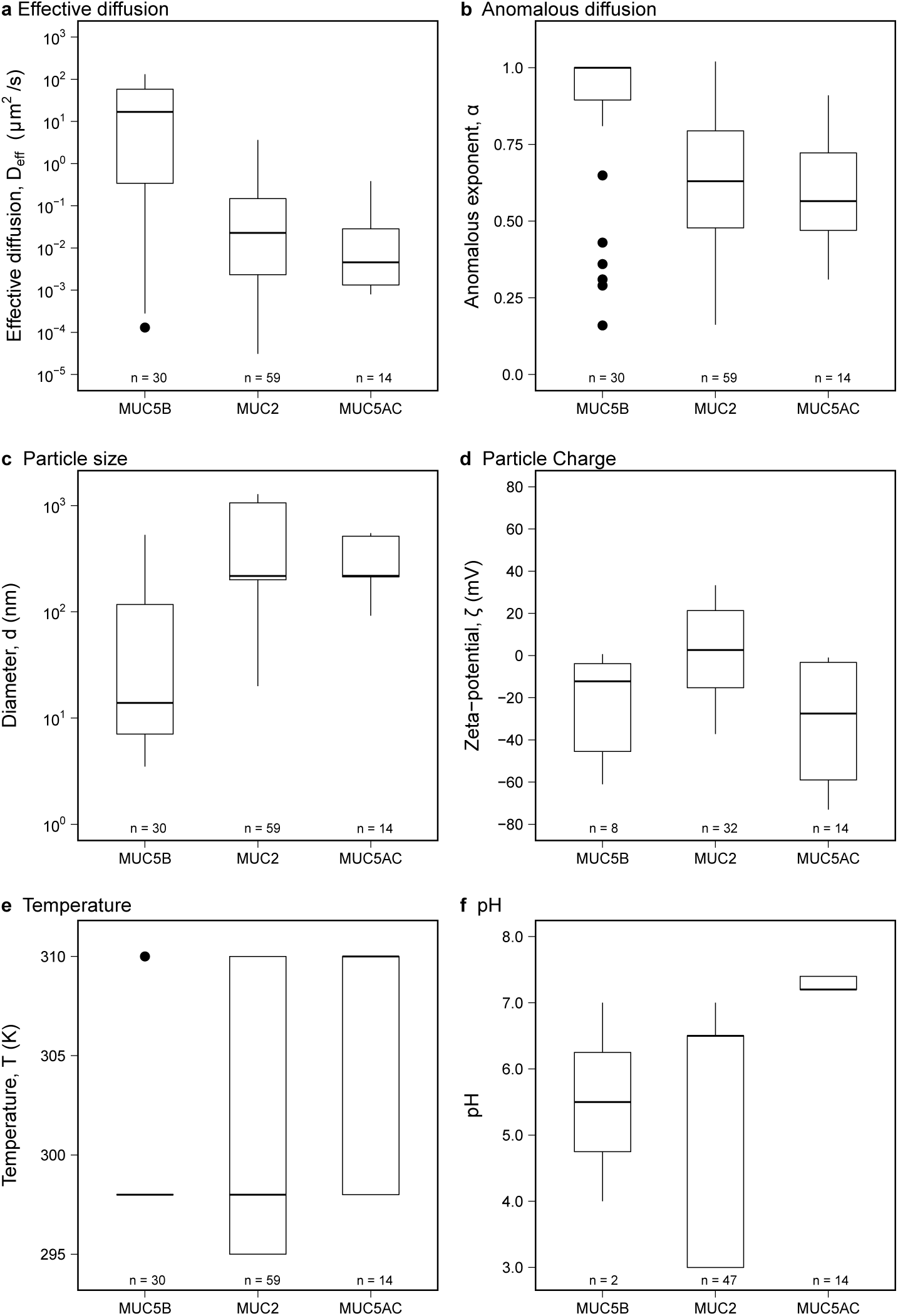
Impact of dominant mucin genes on particle transport. **a**, effective diffusion constant at one second based on dominate mucin. **b**, anomalous exponent based on dominate mucin. **c**, particle size based on dominate mucin. **d**, particle net charge based on dominate mucin. **e**, mucus temperature based on dominate mucin. **f**, mucus pH based on dominate mucin. **a-f**, box plots are ranked by effective diffusion from high to low. The total amount of data points for each dominate mucin is designated as n aligned with their respected dominate mucin for each panel.

**Figure S.4:**
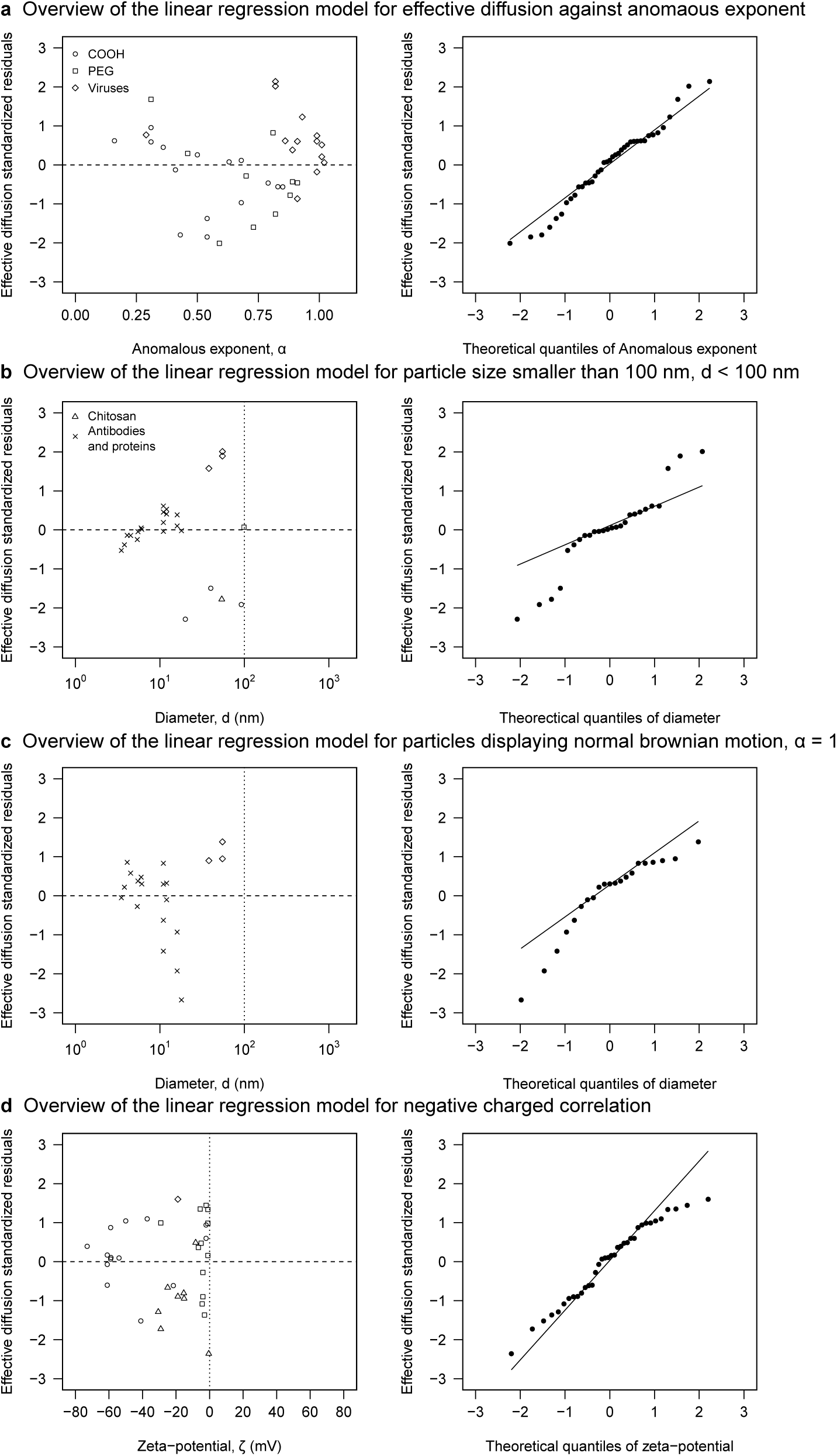
In-depth overview of significant linear regression models. **a**, standardized residual and normal probability of standardized residuals for effective diffusion as a function of anomalous exponent. **b**, standardized residual and normal probability of standardized residuals for effective diffusion as a function of particle size for sizes less than 100 nm. Dotted line is a visualization marker for particles smaller than 100 nm. **c**, standardized residual and normal probability of standardized residuals for effective diffusion as a function of particle size for particles displaying normal brownian motion. Dotted line is a visualization marker for particles smaller than 100 nm. **d**, standardized residual and normal probability of standardized residuals for effective diffusion as a function of zet potential for negatively charged particles. Dotted line is a visualization marker for negatively charged particles. **a-d**, different particle types with corresponding symbols are designated in **a** and **b**’s legend.

